# Assembling a corpus of phosphoproteomic annotations using ProtMapper to normalize site information from databases and text mining

**DOI:** 10.1101/822668

**Authors:** John A. Bachman, Peter K. Sorger, Benjamin M. Gyori

**Affiliations:** Laboratory of Systems Pharmacology, Harvard Medical School, 200 Longwood Avenue, Boston, MA 02115

## Abstract

Protein phosphorylation regulates numerous cellular processes and is highly studied in biology.However, the analysis of phosphoproteomic datasets remains challenging due to limited information on upstream regulators of phosphosites, which is fragmented across multiple curated databases and unstructured literature. When aggregating information on phosphosites from six databases and three text mining systems, we found that a substantial proportion of phosphosites were mentioned at residue positions not matching the reference sequence. These errors were often attributable to the use of residue numbers from non-canonical protein isoforms, mouse or rat proteins, or post-translationally processed proteins. Non-canonical site numbering is also prevalent in mass spectrometry datasets from large-scale efforts such as the Clinical Proteomic Tumor Analysis Consortium (CPTAC). To address these issues, we developed ProtMapper, an open-source Python tool that automatically normalizes site positions to human protein reference sequences. We used ProtMapper coupled with the INDRA knowledge assembly system to create a corpus of 37,028 regulatory annotations for 16,332 sites – to our knowledge, the most comprehensive corpus of literature-derived information about phosphosite regulation currently available. This work highlights how automated phosphosite normalization coupled to text mining and knowledge assembly allows researchers to leverage phosphosite information that exists within the scientific literature.

## Introduction

Protein post-translational modifications (PTMs), phosphorylation in particular, regulate diverse cell functions such as signal transduction and cell fate determination (Pawson & Scott, 2005). Protein phosphorylation has been widely studied and is the focus of sophisticated curated resources such as PhosphoSitePlus (Hornbeck et al., 2012). Misregulation of PTMs plays a prominent role in cancer, which has motivated the generation of proteome-scale datasets such as those from the Clinical Proteomic Tumor Analysis Consortium (CPTAC; https://proteomics.cancer.gov/programs/cptac) (NCI CPTAC et al., 2016; H. Zhang et al., 2016). CPTAC provides collections of ‘omic data, including mass spectrometry-based phosphorylation data, for many proteins within tumors. This data may hold valuable biological insights that could advance cancer diagnosis and treatment, but analyzing such ‘omic datasets can be challenging.

Functional annotations of proteins are critical for generating mechanistic biological insights from ‘omic datasets. These annotations describe key information, such as the identity of kinases that phosphorylate a specific protein site (also known as a phosphosite) and how phosphorylation on a specific site affects protein function. In addition to PhosphoSitePlus, databases such as SIGNOR (Perfetto et al., 2016), and Reactome (Fabregat et al., 2018) summarize much of the available information on phosphosites based on human curation of the literature. However, the proportion of experimentally observed phosphosites with one or more functional annotations is very low, estimated to be ~3% for regulatory annotations and below 3% for functional effects (Needham et al., 2019). Thus, new approaches are needed to capture published information on the function and regulation of phosphosites and to improve the use of information already in databases.

Automated text mining can extend the scope of curated databases by extracting structured information on phosphosites and their regulation from the primary literature (Torii et al., 2015; Valenzuela-Escárcega et al., 2018). However, we show in this paper that inconsistencies in the way phosphosite positions are recorded across research articles, curated databases, and phosphoproteomic datasets makes aggregating phosphosite data challenging. In many cases, phosphosites faithfully extracted by human curators or text mining systems from primary sources cannot be matched to protein reference sequences. Most inconsistencies can be traced back to the use of site positions from nonhuman species, non-canonical isoforms, or post-translationally cleaved proteins. This problem is compounded by mass spectrometry data processing pipelines that map peptide sequences present in multiple isoforms of a given protein to the database identifiers for a non-canonical isoform of that protein (despite the peptide existing in *both* the canonical sequence and the isoform). As a result, many PTMs in mass spectrometry datasets do not match known site positions in databases or literature. Together, these errors cause many known functional annotations to be missed; this exacerbates the already poor coverage of phosphosite annotations and potentially impacts the biological interpretation of phosphoproteomic experiments.

We have developed two resources to help address these problems: 1) an open source software tool, ProtMapper, that resolves inconsistencies among phosphosite annotations and experimentally identified sites, and 2) a large corpus of regulatory phosphosite annotations aggregated from six databases and three text mining tools using the Integrated Network and Dynamical Reasoning Assembler (INDRA) (Bachman et al., 2022; Gyori et al., 2017), which we have linked to ProtMapper. Normalizing sites with ProtMapper resolves inconsistencies between distinct sources by mapping phosphosites from different protein isoforms to known phosphorylation sites in the UniProt human reference sequence (whenever possible). This makes it possible to find annotations for previously undescribed phosphosite positions and to correctly assemble annotations for known positions. Our final corpus contains 37,028 regulatory annotations for 16,332 distinct sites, a 2.6-fold increase over PhosphoSitePlus, which currently represents the most comprehensive database available on protein phosphorylation. Although the assembled ProtMapper corpus is not as extensively curated as the gold-standard PhosphoSitePlus, we show that it is biologically useful: the aggregated and normalized data nearly doubles the number of annotated sites in CPTAC Breast Cancer and Ovarian Cancer datasets, and about half of this increase is due to the ability of ProtMapper to map phosphosite annotations and experimental peptides to the correct reference sequences. These tools allow users to uncover connections across existing bodies of research with the potential to increase understanding of normal and disease biology.

ProtMapper is available under an open-source BSD 2-clause license at https://github.com/indralab/protmapper, and the corpus of phosphosite annotations is available as Supplementary Data with this paper under a CC-BY-NC-SA license. All results from the paper are reproducible from code available at https://github.com/indralab/protmapper_paper.

## Results

### Pathway databases and literature contain annotations of human PTMs that do not match reference sequences

To construct a corpus of phosphosite annotations, we used INDRA to aggregate information from multiple curated databases and text mining systems (**Figure 1A**). Specifically, we included the databases PhosphoSitePlus (Hornbeck et al., 2012), SIGNOR (Perfetto et al., 2016), HPRD (Mishra, 2006), NCI-PID (Schaefer et al., 2009), Reactome (Fabregat et al., 2018), and the BEL Large Corpus (https://biological-expression-language.github.io/), as well as extractions from three text-mining systems: Reach (Valenzuela-Escárcega et al., 2018), Sparser (McDonald et al., 2016), and RLIMS-P (Torii et al., 2015). Each text mining system processed information from a text corpus that contained abstracts and full-text articles (see Methods). From each source we obtained phosphorylation information that included a residue and position on a target substrate. We also extracted available information on upstream regulators (e.g. the relevant kinase), downstream effects, and the phosphosite function, the latter primarily in the form of preconditions for a protein to participate in a downstream reaction (e.g., MAPK1 *phosphorylated at T185 and Y187* phosphorylates RPS6KA1).

**Figure 1.**
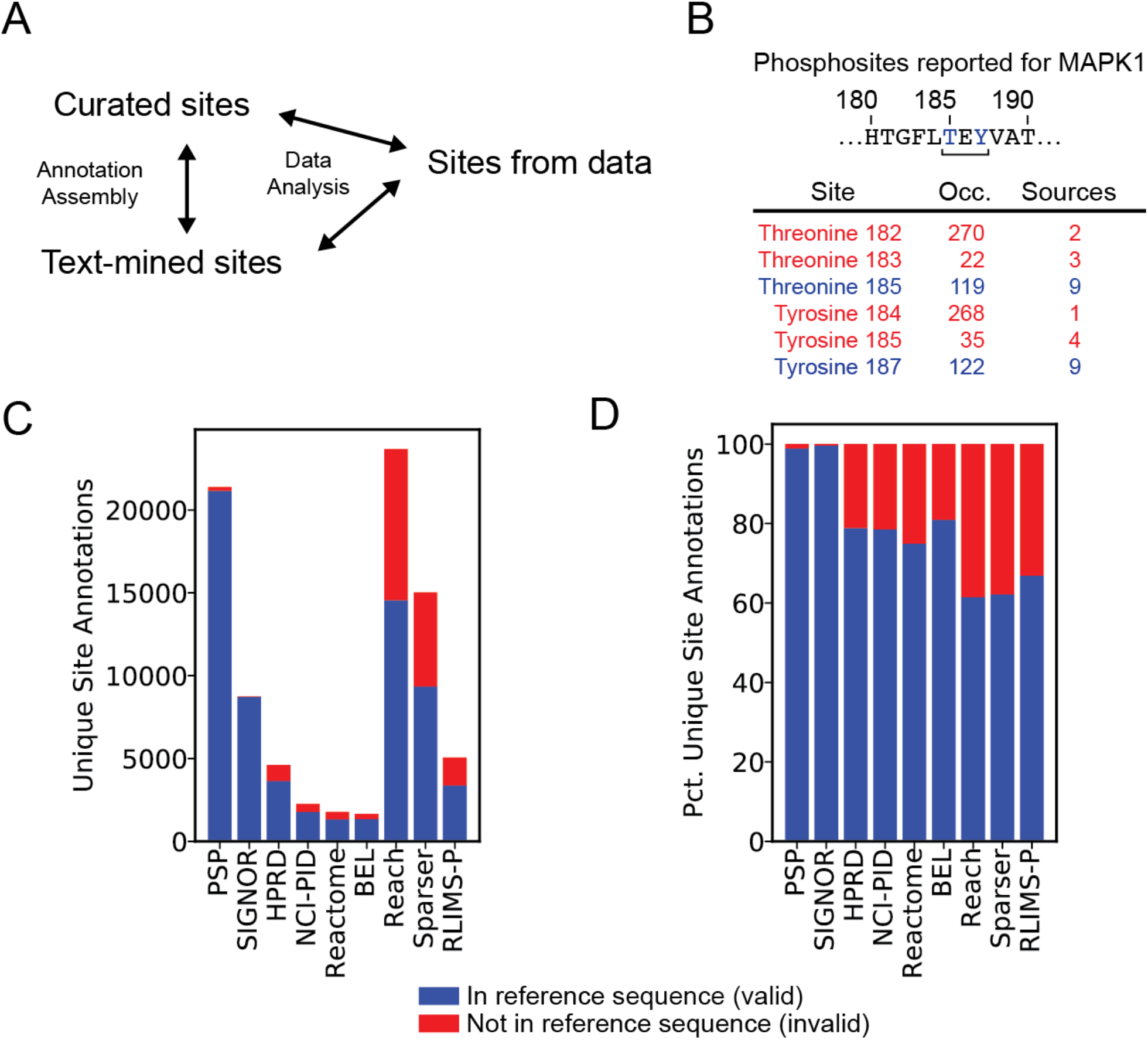
Inconsistencies in reported positions of phosphosites. **(A**) Interpreting phosphoproteomic data requires linking experimentally observed phosphosites to annotations assembled from both curated databases and literature. **(B**) An example of inconsistent site positions for the T/Y activation motif in human MAPK1. Relevant sequence of the canonical isoform from UniProt is shown above with T and Y residues shown in blue. “Occ.” denotes the total number of occurrences (reactions or sentences) across all sources; “Sources” denotes the number of sources reporting the site at the given position. Non-canonical positions are shown in red; canonical positions in blue. **(C**) Number of unique site annotations found in human reference sequence, by source. A site annotation consists of a unique combination of regulator, substrate, residue, position. **(D**) Percentage of unique site annotations matching human reference sequence, by source.

We next sought to evaluate the quality of the assembled corpus of phosphosite annotations. To do this, we leveraged the UniProt (The UniProt Consortium, 2019) and NCBI RefSeq Protein (O’Leary et al., 2016) databases, which catalog sequences for all protein isoforms and distinguish between a protein’s reference isoform and its less common splice variants. We found that PTMs in the corpus were frequently listed at positions that did not contain a phosphorylatable residue (serine, threonine, or tyrosine) in the corresponding reference sequence; this issue has also been observed previously by others (Babur et al., 2021; Huckstep et al., 2021). For example, in human MAPK1 (mitogen-activated protein kinase 1, also known as ERK2), the positions of the T and Y residues in the T-X-Y activation motif are recorded across the literature in a variety of amino acid positions including 183/185, 184/186, and 185/187. However, the UniProt and NCBI RefSeq reference sequences list the T-X-Y motif as occurring at residues 185-187 (**Figure 1B**). In this case, the disparities in position prevented accurate aggregation of database annotations and literature-based evidence on the phosphorylation of MAPK1, an event essential for cell proliferation (**Figure 1B**). Thus, even minor phosphosite position shifts between sources can confound understanding of important biological phosphorylation events.

To quantify the extent of site annotation inconsistencies we performed a systematic assessment of the assembled phosphorylation corpus. In this paper we refer to a site annotation as having a “valid” position when the residue listed in the annotation (i.e., S, T, or Y) matches the residue in the reference sequence at the annotated position; otherwise the annotated position is “invalid.” According to this definition, we found that the total number of unique site annotations and the fraction of annotations that were valid varied from 62% to over 99% across sources (**Figures 1C, 1D**; **Table 1**). Among databases, PhosphoSitePlus had by far the largest number of unique site annotations (21,390) (**Figure 1C**, **Table 1**), and the second-highest proportion (98.9%) of valid site annotations (**Figure 1D, Table 1**). This likely reflects the fact that PhosphoSitePlus is actively maintained by a team of expert curators. Text mining contributed a large number of site annotations, with both Reach and Sparser independently extracting 23,666 and 15,030 unique site annotations respectively – more than any source except PhosphoSitePlus (**Figure 1C; Table 1**). Notably, over 35% of the text-mined annotations were invalid, but it was not clear *a priori* whether the problem lay with machine reading errors or technically correct extractions referring to non-reference protein sequences (**Figure 1D, Table 1**). Remarkably, ~22-25% of the sites in the NCI Nature Pathway Interaction Database (PID) and Reactome, two widely used human-curated pathway databases, also were not valid (**Figure 1D, Table 1**).

**Table 1.**
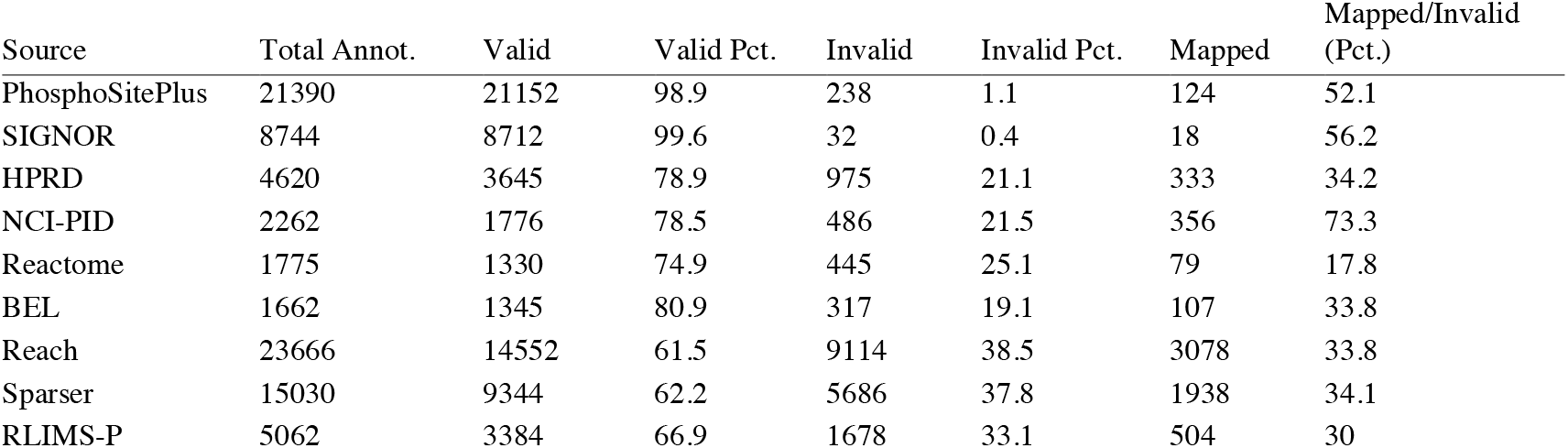
Annotations linked to valid, invalid, and mappable sites, by source.

### Site inconsistencies can be traced to the original literature as well as curation and text mining errors

To address the problem of invalid phosphosite annotations we investigated why they occur. By reviewing extracted site information, we found that many invalid site positions were caused by inconsistent numbering of site positions in the original source literature. These non-canonical positions were then propagated to phosphosite annotation databases by human curators and text mining systems (Table 2). We identified four common reasons for the invalid phosphosite positions found in primary research papers: (i) use of mouse or rat protein sequences when describing human PTMs, (ii) use of non-canonical protein isoforms generated by alternative splicing, (iii) use of sequence positions from a different member of a multi-gene family, and (iv) differences in residue number arising from proteins that undergo post-translational modifications such as the cleavage of a signal peptide or initial methionine (**Table 2**).

**Table 2.** Types of inconsistencies for invalid sites in curated databases and text mining results

| Type of inconsistency | Example Site | Description |
| --- | --- | --- |
| Site position from other species | BAD S112 | Mouse site, corresponds to human S75 |
| Site position from other isoform | RPS6KB1 T389 | Position from isoform Alpha II |
| Site position from other family member | RPS6KA6 S221 | Position from RPS6KA1 |
| Initial methionine cleavage in mature protein | LCK Y393 | Y394 in reference sequence |
| Signal peptide cleavage in mature protein | EGFR Y1173 | Position after cleavage of initial 24aa |
| Curator error: wrong residue | BRAF T151 | Should be BRAF S151 |
| Curator error: site from wrong protein | JAK2 Y485 | Site on EPOR where JAK2 binds |
| Text mining error: misidentified protein | HNRNPU S268 | Ambiguous “p120” in sentence refers to CTNND1 |
| Text mining error: site from wrong protein | IRS1 T308 | Site for AKT1, mentioned in same sentence |
| Text mining error: misidentified site | TP53 S1 | “Text S1” extracted as site |

In addition, we identified some examples of curator and machine-reading errors. Curator errors included misannotation of a serine site as a threonine and incorrect annotation of a site that is actually on an interacting or closely related protein. In the latter case, NCI-PID listed a phosphorylation of JAK2 kinase at position Y485, but the Y485 phosphosite actually belongs to erythropoietin receptor precursor (EPOR), a protein that binds JAK2 (**Table 2**). Text mining systems made a variety of errors, such as incorrectly identifying gene and protein names (incorrect grounding), which caused phosphosites to be assigned to the wrong protein. For example, in the sentence *“p120’s phosphorylation at S268 was found to contribute to cellular foci formation”* (Hong et al., 2016), the ambiguous label “p120” was misidentified as gene *HNRNPU*, but it is clear from the context of the paper that the authors were referring to the *CTNND1* gene, which is colloquially known as “p120” in some fields. In other cases, machine readers linked sites to the wrong protein in a sentence. For example, in the phrase *“phosphorylation of IRS-1 and subsequent activation Akt at Thr308” (sic)* (Leclerc et al., 2010), the site T308 was incorrectly associated with IRS1 rather than AKT. A third type of error resulted from the misidentification of site-like text, for example a (non-existent) phosphoserine site TP53 S1 was extracted from the text “…*as determined by p53 stabilization and phosphorylation (see Figure S3 and Text S1)”* (Coppé et al., 2008) (**Table 2**). Collectively, these site errors interfere with the generation of accurate, well-annotated phosphosite resources and phosphoproteome networks necessary for the analysis of mass spectrometry data (**Figure 1A**). The problem is greater for machine-reading systems, which struggle to understand context, but even human curators make errors.

### Normalizing phosphosite information by mapping sites to human reference sequences

Having identified common causes of invalid phosphosite annotations, we sought an approach to account for inconsistencies and unify information across curated databases, text mining, and experimental data (“site normalization”). To do this, we developed ProtMapper, a systematic method to robustly normalize invalid, non-canonical phosphosites to their corresponding valid positions in human reference sequences wherever possible (**Figure 2A**). ProtMapper is implemented as open-source Python software with source code available at https://github.com/indralab/protmapper. In addition to performing site normalization, ProtMapper includes useful tools for analyzing proteomic data, such as a wrapper around PhosphoSitePlus site information, mappings between UniProt and NCBI RefSeq Protein identifiers and gene symbols, and sequence lookup for both NCBI RefSeq Protein and UniProt (**Figure 2B**).

**Figure 2.**
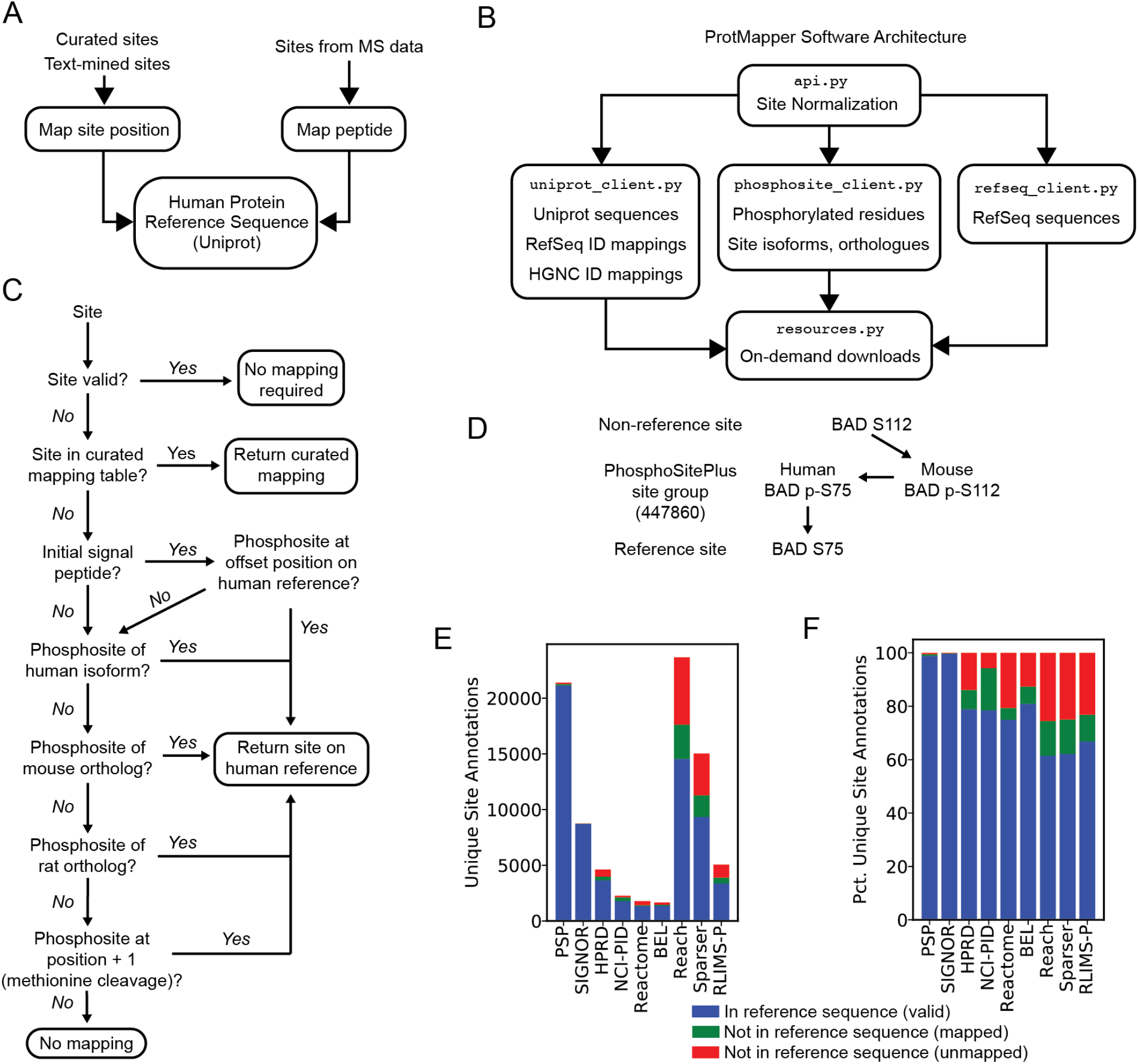
Mapping invalid phosphosites to human reference sequence positions. **(A**) The ProtMapper maps identifiers and site positions for phosphosite annotations experimental phosphopeptides to corresponding positions on the human UniProt reference sequence. **(B**) ProtMapper Software Architecture. Site normalization is implemented in the api.py module, which draws on additional modules for ID normalization, protein sequences, and phosphosite information. Resource files are downloaded as needed at run time using functions in resources.py. **(C**) Site normalization procedure. A site, including residue and position, is checked for validity against the UniProt reference sequence. If invalid, a series of mappings are attempted, starting with a curated mapping table and proceeding through positions for known phosphorylation sites after signal peptide cleavage (if any); known phosphosite positions from human isoforms and mouse and rat orthologues; and known phosphosites at position + 1, a common inconsistency due to the cleavage of the initial methionine of the protein. **(D**) PhosphoSitePlus site groups. If a known phosphorylation site with the given position is found on an alternative sequence (e.g., the mouse ortholog), it can be mapped back to the human reference position via PhosphoSitePlus site groups, which link together corresponding sites from human isoforms and nonhuman orthologs. **(E**) Counts of valid, mappable, and invalid/unmappable site annotations, by source. **(F**) Percentages of valid, mappable, and invalid/unmappable site annotations, by source.

The goal of the ProtMapper site normalization tool is to determine the most likely reference position (if one exists) for reported PTM sites that do not match a protein’s reference sequence. To accomplish this, ProtMapper first determines whether the site is valid with respect to the reference sequence; if it is, no mapping is required (**Figure 2C**). For reported sites that are invalid with respect to the reference sequence, ProtMapper first compares the site to a mapping resource that we generated by human curation. We developed this resource by reviewing frequently occurring non-canonical site positions in NCI-PID (a resource with a high proportion of invalid sites, **Figure 1D**) and manually curating 134 mappings to appropriate valid site positions. ProtMapper compares invalid site positions against this resource first, and if the site is in the table, ProtMapper returns the curated mapping position. This step is intended to deal with the most frequently encountered invalid sites; as an open-source resource, the curated mappings can also be expanded over time.

Second, if a site is not found in this mapping table, ProtMapper uses annotations in Uniprot to determine whether the protein contains a signal peptide that, when cleaved off, could explain a shift in phosphosite numbering; signal peptides are commonly found on cell surface receptors. If ProtMapper finds a matching residue on the reference sequence when the invalid position is shifted by the length of the signal peptide, then it returns the shifted position (**Figure 2B**).

Third, ProtMapper determines whether the invalid position is a *known* site of phosphorylation on a closely related protein sequence in human isoforms and mouse, and rat orthologues (**Figure 2B**). To determine the set of related proteins on which a valid phosphosite might exist, ProtMapper draws on data from PhosphoSitePlus *site groups*, which catalogue corresponding positions between homologous sequences (human and non-human proteins and isoforms), allowing positions of known phosphorylation sites to be mapped between sequences without additional sequence alignment. For example, the (invalid) site human pBAD-S112 can be mapped correctly to human pBAD-S75 because murine pBAD-S112 and human pBAD-S75 represent the same site group (**Figure 2C**). The set of related proteins are tested sequentially in a fall-through fashion with the most conservative options considered first. Thus, if an invalid site position can be associated with a known phosphorylation site on a different human isoform, ProtMapper returns the corresponding reference position without considering mouse or rat orthologs (**Figure 2B**). Note that PhosphoSitePlus site groups do not include corresponding gene positions among paralogs, so mappings between gene family members are not automatically handled by this procedure (e.g., mapping the activating site T308 on the kinase AKT1 to the corresponding T309 on AKT2). However, we find these types of inconsistencies to be much less common than those involving alternative human isoforms or non-human species.

In a final step, the ProtMapper checks for “off-by-one” shifts in position due to the use of sequences that exclude a post-translationally cleaved initial methionine residue. For example, if Y10 is an invalid position with respect to the target reference sequence, ProtMapper checks whether the reference sequence has a matching residue (i.e., Y) at the position one greater (i.e, 11) that is known to be phosphorylated in PhosphoSitePlus; if so, the updated position is assigned (**Figure 2C**).

When we applied ProtMapper to our corpus of phosphosite annotations (**Table 1**), we were able to identify reference human positions for many invalid site annotations in the corpus (**Figures 2D** and **E**, **Table 1**). Among the curated databases examined, NCI-PID had the largest fraction of initially invalid site annotations that could subsequently be mapped to a canonical residue (73% “mappable” site annotations). In absolute numbers, text mining systems had by far the largest number of mappable invalid site annotations. ProtMapper identified valid reference sequence positions for a total of 5,520 sites out of the total 16,478 invalid site annotations that had been extracted by three different reading systems (33%; **Table 1**). Across the three text mining systems, the percentage of mapped sites was very similar, with 34%, 34%, and 30% of invalid annotations mapped for Reach, Sparser, and RLIMS-P respectively (**Table 1,** “Mapped/Invalid (Pct)”). Thus, site normalization with ProtMapper increased the proportion of valid text mined site annotations by 15-21%, depending on the text mining system (**Table 1**, “Mapped” divided by “Valid”).

### Accuracy of automatically inferred mappings for literature-derived sites

The approach that ProtMapper takes to normalize invalid phosphosites has the potential to miss legitimate mappings to valid positions (i.e., false negative mapping) and to erroneously associate invalid sites with reference sequence positions (i.e., false positive mapping). This is a particular problem for normalizing text-mined sites due to the high frequency of technical errors (**Table 2**). To determine the accuracy of ProtMapper when applied to sites mined from the literature, we manually reviewed a random sample of 100 invalid text-mined phosphosites. It was feasible to perform this analysis on only a sample of sites because evaluating the correctness of site mappings requires careful inspection of the original source literature (the precise criteria we used to evaluate the accuracy of site mappings are described in Methods). The dataset containing the phosphosites, ProtMapper mappings, and the assessments of correctness is available at https://github.com/indralab/protmapper_paper.

Overall, manual curation of ProtMapper results yielded estimates of 95% precision and 74% recall, with the high precision figure due to 41 of 43 technical reading errors being correctly ignored (**Table 3**). Thus, ProtMapper is robust to both false positive and false negative errors. Of the 100 invalid text-mined phosphosites in our sample, 43 were caused by text mining errors of the types described in **Table 2**. A spurious mapping of these incorrectly extracted sites to a reference sequence would therefore represent a false positive, but we only observed two such errors. For example, in the sentence “*phosphorylation of IRS-1 and subsequent activation Akt at Thr308*” (*sic*), the site T308 was incorrectly associated with IRS1 due to a reading error (**Table 2**; row 52 in curation dataset). There is no threonine at position 308 on IRS1, but there is a threonine at position 309 that is known to be phosphorylated based on PhosphoSitePlus. ProtMapper incorrectly associated “Thr308” with IRS1-T309 under the assumption that it is an off-by-one inconsistency arising from cleavage of the initiator methionine. Only one other reading error resulted in an erroneous mapping to a human reference sequence (row 62 in the curation dataset). Since both false positive annotations resulted from the “off-by-one” rule (**Figure 2C**), we tried inactivating the “off-by-one” rule and found that this increased ProtMapper’s precision to 100% but reduced recall to 68% (decreasing the F1-score to 0.81). As a compromise, we made inactivation of the “off-by-one” rule optional in ProtMapper, making it possible to tune the precision/recall tradeoff for different use cases. In the case of the remaining 41 text mining errors, phosphosites were not mapped to a canonical sequence by ProtMapper (**Table 3**). Importantly, this information from ProtMapper—i.e., that an invalid site position cannot be mapped to a valid one—can be used to filter out site annotations, eliminating these text mining errors from downstream usage.

**Table 3.** Summary of results for mapping invalid sites extracted from the literature using machine reading, with a sample size of 100 sites.

| Category | ProtMapper mapped the site | ProtMapper did not map the site | Total |
| --- | --- | --- | --- |
| Machine reading extraction correct | 42 (true positive) | 15 (false negative) | 57 |
| Machine reading extraction incorrect | 2 (false positive) | 41 (true negative) | 43 |
| Total | 44 | 56 | 100 |
| Precision |  |  | 0.95 |
| Recall |  |  | 0.74 |
| F1 |  |  | 0.83 |

The remaining 57 invalid sites were correctly extracted from the source literature by the text mining systems (i.e., the extracted sites faithfully represented the underlying text). Among these, ProtMapper was able to map 42 sites (74%) to correct reference sequence positions (true positive mappings). To determine which mapping rules (**Figure 2C**) were used most often for these sites we grouped curation results for the 100 sites described above according to the type of mapping (**Table 4**). We found that all mapping types (shown in **Figure 2C**) were invoked multiple times in our sample; mapping a mouse site position to the corresponding human site was the most commonly applied mapping rule. The “off-by-one” inferred methionine cleavage rule was the only rule that led to incorrect mappings, as noted above (**Table 3**).

**Table 4.** Total and correct site mappings of different types in the curation dataset among the 44 sites that were mapped by ProtMapper.

| Mapping type | Number (total) | Number (correct) |
| --- | --- | --- |
| Inferred mouse site | 13 | 13 |
| Manually curated mapping | 10 | 10 |
| Inferred methionine cleavage | 7 | 5 |
| Reference mismatch between UniProt and PSP | 6 | 6 |
| Inferred alternative isoform | 4 | 4 |
| Inferred signal peptide cleavage | 4 | 4 |

A total of 15 of 57 invalid sites (26%) were correctly extracted from text but could not be associated with a corresponding reference sequence by ProtMapper; these represent false negatives (**Table 3**). We re-read the original papers used by the text mining systems to understand why ProtMapper was unable to normalize these sites. In six cases we found that the site referenced a position in a related human isoform, mouse or rat protein, or methionine-cleaved sequence that was not known to be phosphorylated in PhosphoSitePlus, so no mapping was made. This is not unexpected given that even a large database such as PhosphoSitePlus doesn’t have a complete catalog of all possible human, mouse and rat phosphosites. Two additional sites involved orthologous proteins from Drosophila and Xenopus, organisms that are not generally included in PhosphoSitePlus (for example site S381 on *Xenopus* Msi2, row 61 in the curated dataset). In one case, the invalid site was due to an error in the primary article itself: a reference to tyrosine 828 was listed as “T828” rather than Y828 (row 77 in the curated dataset). In the remaining six cases it was not possible for a human curator to identify a reason for the sequence position discrepancy.

Though it is possible in principle, we did not encounter any examples of mappings in which an invalid site was extracted correctly by text mining and then mapped to a reference position that was inconsistent with the original article’s context. For example, with a different precedence of mapping rules (**Figure 2C**), a site referring to a position in an alternative human protein isoform might have been mapped to a different (incorrect) position corresponding to a known mouse phosphosite; however, we observed no errors of this type.

### Text mining tools identify many uncurated regulators of phosphorylation

After combining PTM annotations from databases and literature using INDRA and normalizing the data with ProtMapper (**Figure 2**), we were left with a corpus of 37,028 regulatory annotations for 16,332 human sites (the regulatory annotations along with the underlying sentences for text mined annotations are available as Supplementary Data). Our combined, normalized corpus contains 2.6 times as many regulatory annotations (37,028 vs. 13,989) and 1.7 times as many distinct sites (16,332 vs. 9,553) as PhosphoSitePlus itself. We measured the specific contributions of text mining and databases to the normalized dataset by comparing the overlap of regulator-site pairs between (i) PhosphoSitePlus (the largest curated database), (ii) other widely used pathway databases (HPRD, Signor, NCI-PID, Reactome, and the BEL Large Corpus), and (iii) the aggregated output of the Reach, Sparser, and RLIMS-P text mining systems. We made these measurements after normalization by ProtMapper, thereby excluding invalid and unmappable sites for greater precision. A visual representation of this overlap is shown in **Figure 3**.

**Figure 3.**
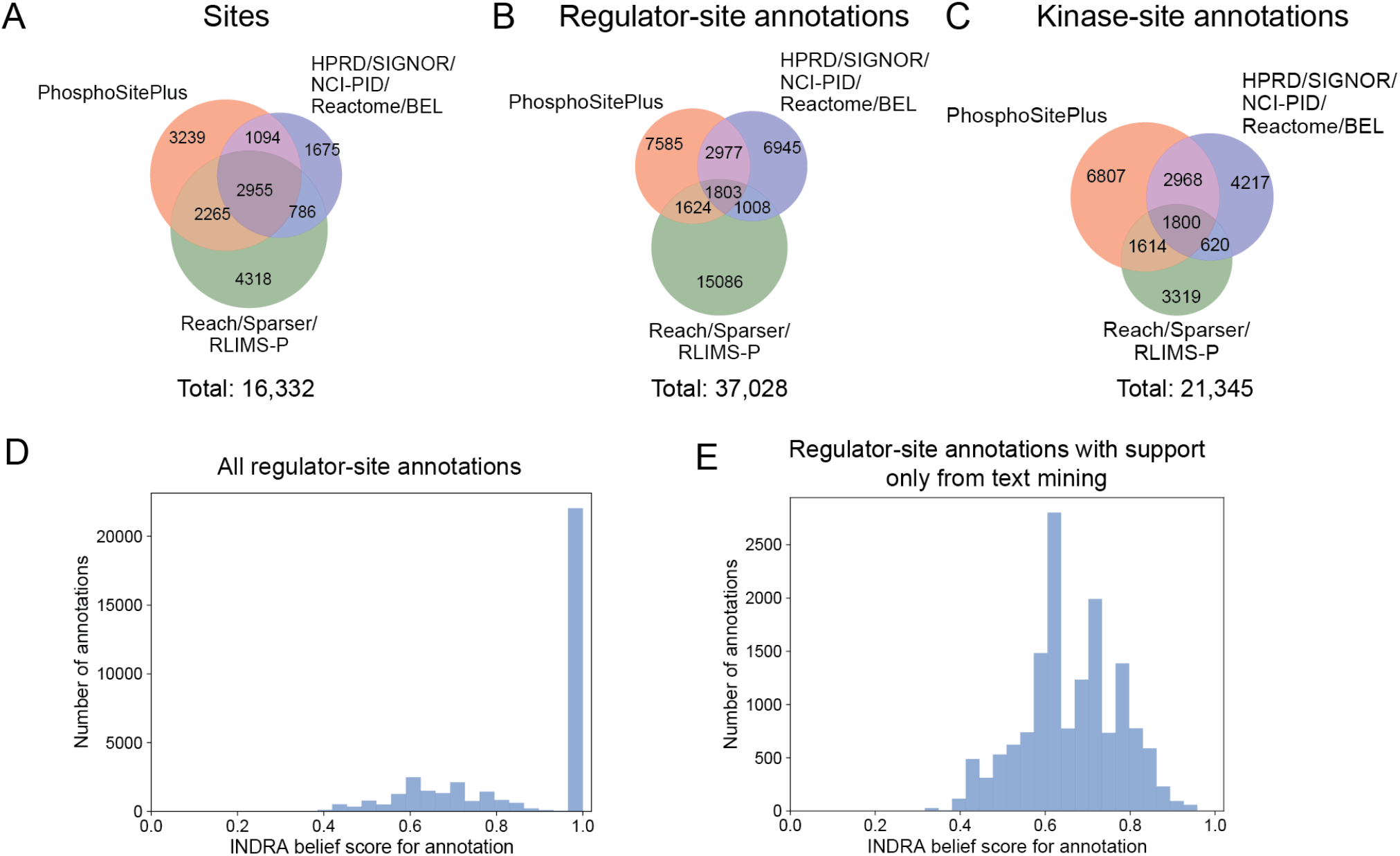
Overlap of phosphosite information between PhosphoSitePlus, the combined output of the HPRD, SIGNOR, PID, Reactome, and BEL databases, and the combined output of the Reach, Sparser and RLIMS-P machine reading systems. **(A**) Venn diagram of unique sites (irrespective of regulator) reported by PhosphoSitePlus, other pathway databases, and machine reading systems. **(B**) Venn diagram of unique regulator-site pairs reported by PhosphoSitePlus, other pathway databases, and machine reading systems. **(C**) Venn diagram of unique regulator-site pairs filtered to human-only kinases and removing protein families and complexes, reported by PhosphoSitePlus, other pathway databases, and machine reading systems. **(D**) Histogram of belief scores for each regulator-site annotation, calculated by INDRA. Belief scores for all regulator-site annotations are shown, including ones with direct or indirect curated database support (belief set to 1). **(E**) Histogram of belief scores for the subset of regulator-site annotations that only have support from text mining, calculated by INDRA.

We found that text mining contributed a large proportion of new annotations to the corpus. Specifically, text mining extracted one or more upstream regulators for 10,324 sites, of which 5,104 (49%) did not have machine-readable regulatory annotations in PhosphoSitePlus, and 4,318 (42%) did not have regulatory annotations in any curated databases (**Figure 3A**). Text mining also identified 19,521 unique regulator-site pairs, 15,086 of which were absent from all curated databases (**Figure 3B**). While the union of all regulator-site annotations obtained from Reach, Sparser, and RLIMS-P was fairly large (19,521; **Figure 3B**), only 1,371 (7%) of these were extracted by all three, highlighting the value of combining results from multiple systems.

In comparing text mined and curated annotations, certain key differences must be taken into account. First, in addition to direct interactions between kinase and substrate, text mining also extracts information about kinases that lie further upstream from the target as well as non-kinase regulators (e.g., growth factors); this expands the corpus of interactions relative what has been previously curated. Moreover, while curated databases generally limit phosphosite annotations to specific kinases, Reach and Sparser extract and normalize phosphorylation events in terms of kinase classes using FamPlex identifiers, a taxonomy of protein families and complexes (e.g., “ERK”, “AKT”, “AMPK”) (Bachman et al., 2018). The inclusion of annotations at the family level is useful because many observations of phosphosite regulation (e.g., studies using poly-selective kinase inhibitors) do not distinguish between individual members (e.g., ERK1 vs. ERK2). To estimate the extent to which the larger number of annotations in the normalized corpus reflected the inclusion of these other types of regulators, we repeated the comparison with curated databases by restricting the text mining dataset to human kinases, excluding non-kinase regulators as well as kinase families and complexes. With this restriction in place we still found that text mining contributed a substantial body of new information, with 3,939 unique human kinase-site pairs reported by machine readers that were not yet present in PhosphoSitePlus, a subset of which (3,319) did not appear in any curated database (**Fig 3C**).

A second key difference between curated and text-mined annotations is accuracy. As discussed above (**Tables 2 and 3**), text mining systems can make technical errors in extracting biological interactions, while curation errors are much less common. Because phosphosite annotations in our normalized corpus include only those phosphosites where the site corresponds to the substrate protein many text mining errors are filtered out (**Table 3**); however, incorrect extraction of the regulator protein is also possible. To address this, we used INDRA’s random forest Belief Model (Bachman et al., 2022). to assess the technical reliability of the text-mined annotations in the normalized corpus. INDRA Belief scores are an estimate of each annotation’s probability of being technically correct across the phosphosite regulator, substrate, residue and position. Of the 37,028 annotations, 22,040 had direct or indirect support (here, indirect support means that we consider an annotation on a specific member of a protein family/complex to also support the annotation for the given family/complex, consistent with INDRA’s approach to belief calculation) from at least one curated database and were assigned a belief value of 1 (**Figure 3D**). For the remaining 14,988 annotations with support only from text mining, predicted belief scores ranged from 0.32 to 0.96 with a median score of 0.66 (**Figure 3E**). The belief scores can be used to filter the corpus of annotations for use in downstream data analysis applications to control the false discovery rate.

### ProtMapper normalizes peptides in phosphoproteomic datasets to align with the reference sequence

As a use case for ProtMapper, we used it to normalize and analyze two large-scale phosphoproteomic datasets generated using mass spectrometry (one on breast cancer, the other on ovarian cancer, see Methods) and released by the NCI Clinical Proteomic Tumor Analysis Consortium (CPTAC) (NCI CPTAC et al., 2016; H. Zhang et al., 2016). We found that both protein identifiers and phosphosite sequence positions in the CPTAC data suffered from alignment issues with respect to reference phosphosite annotations. In common with most phosphoproteomic datasets, CPTAC phosphoproteomic data identify phosphosites using NCBI RefSeq Protein IDs and corresponding HGNC gene symbols (Tweedie et al., 2021). However, the database sources from which we assembled phosphosite annotations (with the exception of HPRD) use UniProt IDs to identify phosphorylated proteins. To map between these sources of protein identifiers we implemented RefSeq-to-UniProt and HGNC-to-UniProt mapping procedures in ProtMapper. We observed that a higher proportion of phosphosites could be mapped to a UniProt identifier when we used HGNC rather than RefSeq identifiers as the basis for mapping: < 0.1% of HGNC-to-UniProt mappings failed as compared to 6-8% of RefSeq-to-UniProt mappings. This is due to the fact that HGNC gene IDs do not distinguish between protein isoforms whereas RefSeq identifiers do, and isoform-to-isoform matching between RefSeq and UniProt can fail due to non-overlapping sets of protein sequences contained in the two databases.

After mapping protein identifiers to reference UniProt IDs we found that >22% of sites in CPTAC datasets were reported to lie on non-canonical positions with respect to the reference sequence. When we examined the original RefSeq identifiers for these proteins, we saw that they represented protein isoforms other than the canonical isoform to which the reference sequence corresponds. This presents a problem for analyzing the phosphosite dataset with respect to regulatory annotations: experimentally observed phosphopeptides grounded to isoform-specific identifiers would frequently be considered as having no regulatory annotations in phosphosite databases when in fact there may exist valid annotations associated with the *reference sequence* site position. Of course, phosphoproteomic datasets provide the amino-acid sequence of the peptide on which the given phosphosite was detected, making it possible to determine whether the peptide is unique to a non-canonical isoform or is also present (with identical sequence) on the canonical isoform. In the latter case, the peptide can be matched to the reference sequence allowing the given phosphosite to be mapped to a valid reference sequence position. We implemented a peptide-based mapping function in ProtMapper to normalize such sites and found that in 98% of cases ProtMapper could map phosphosite positions recorded in CPTAC as lying on a non-canonical isoform to a valid position on a reference sequence with the exact same peptide. These findings show that phosphoproteomic data analysis can suffer from the use of non-canonical site positions even when there is not experimental evidence that only the non-canonical isoform is modified. ProtMapper can map these site positions such that they are valid with respect to the reference sequences, making it possible to exploit the extensive annotations in phosphorylation databases.

### Annotation assembly and site normalization increases functional annotations of experimentally observed phosphosites

To determine the impact of site normalization of both regulatory annotations and experimental phosphopeptides, we converted CPTAC HGNC identifiers to UniProt identifiers and performed peptide-based site mapping to reference sequences. We then counted the number of annotated phosphosites in the CPTAC datasets under different analysis conditions to assess both the individual and cumulative contributions of text mining and curated databases (**Table 5**). Overall, we found that combining text mining with annotation site normalization substantially increased the proportion of annotated phosphosites. As expected, adding annotations from text mining systems and curated databases other than PhosphoSitePlus increased the number of annotations over PhosphoSitePlus alone, with or without site normalization (**Table 5**, “All” vs. “PSP-only”). For instance, for the ovarian cancer data set, text mining results more than doubled the number of annotations per site as compared to the databases alone (4.85 vs. 2.17, **Table 5,** row 15 vs. row 14); a combination of text mining with databases yielded 3.73 annotations per site, a 72% increase over databases alone.

**Table 5.**
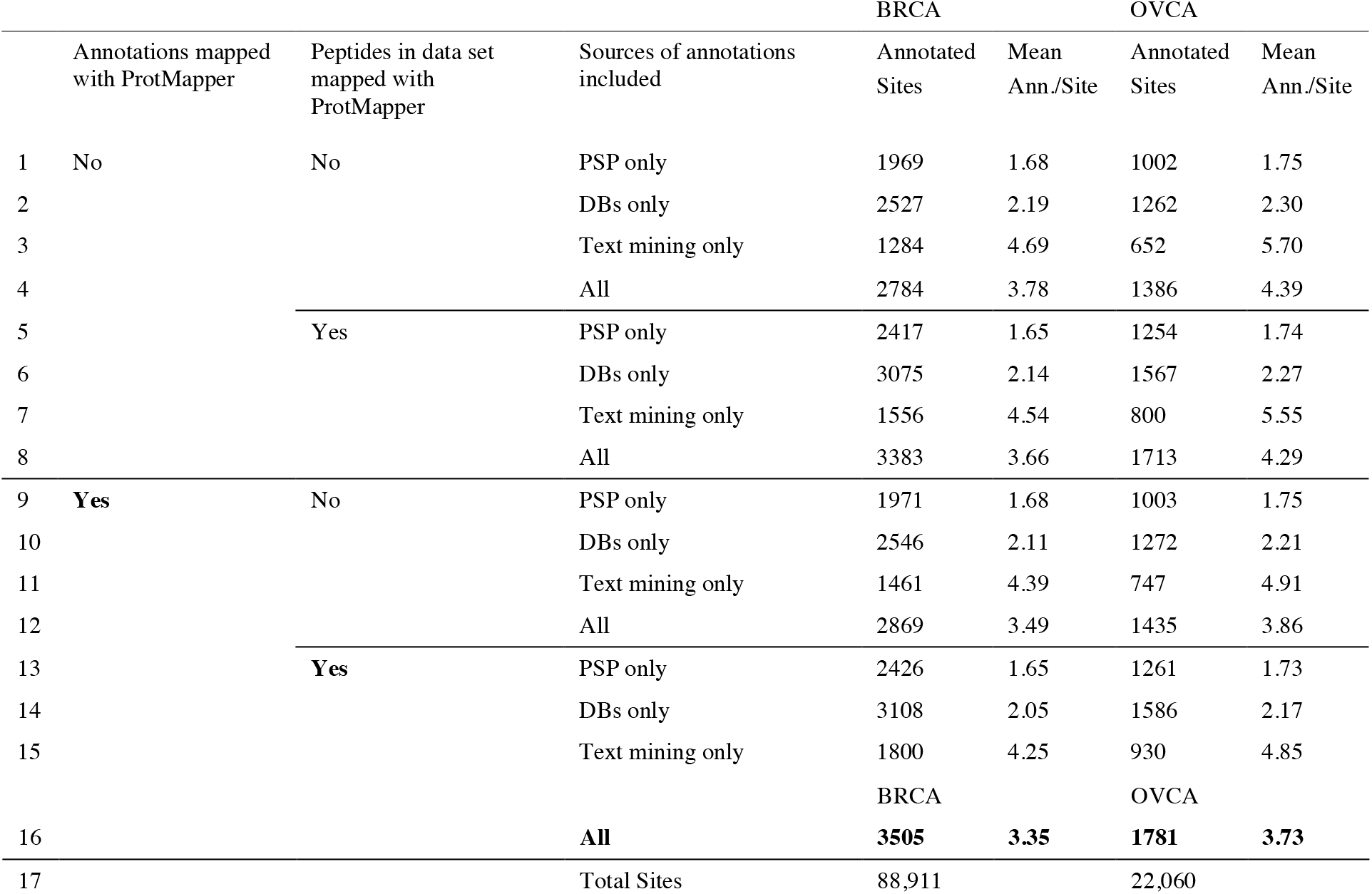
Regulatory annotations of sites in the CPTAC Breast (BRCA) and Ovarian Cancer (OVCA) phosphoproteomic datasets with and without site normalization. PSP only: annotations when using only PhosphoSitePlus, DBs only: annotations when using only curated databases including PhosphoSitePlus, Text mining only: annotations when using only text mining systems, All: annotations when using all sources. The condition in which ProtMapper is applied to both the annotations and the data is shown in bold font.

For the number of unique annotated sites, the inclusion of text mining led to a smaller proportional increase over databases alone: for instance, for the breast cancer data set, the number of annotated sites increased from 3,108 to 3,505 (13%, **Table 5**, row 16 vs. row 14). This shows that text mining systems primarily contributed new regulatory information about sites that already had at least one regulatory annotation in databases. Despite this fact, using the three text mining systems *alone* (i.e., using no human-curated sources) yielded annotations for ~74% of the sites annotated using PhosphoSitePlus (for both breast cancer and ovarian cancer), and included a larger number of annotations per site (**Table 5,** row 15 vs. row 13). This suggests that text mining tools can already meaningfully contribute to human curation efforts by accurately consolidating certain types of information on phosphosites.

We found that site normalization by ProtMapper played a valuable role in data analysis: it increased the number of annotated sites (and annotations per site) regardless of the sources of annotations included. Using all sources of annotations, remapping the peptides in the CPTAC dataset to reference positions increased the annotated sites from 2,784 to 3,383 for breast cancer data, even without mapping site annotations to reference positions (**Table 5,** row 8 vs. row 4). Site mapping further increased the number of annotated sites, reaching a maximum of 3,505 annotated sites for breast cancer data, a 78% increase over the 1,969 sites obtained using PhosphoSitePlus with no ProtMapper normalization (**Table 5**, row 16 vs row 1). Site mapping had a particularly large effect when using only text-mined sources (1,800 vs. 1,556 annotated sites, **Table 5**, row 15 vs. row 7), reflecting the large proportion of text mined information associated with non-canonical site positions. Taken together, these findings with the widely used CPTAC dataset demonstrate the value of normalizing sites for functional annotations and phosphopeptide sequences. By combining site normalization with additional data from text mining and curated databases, we find that ProtMapper can substantially increase the proportion of experimentally observed phosphosites having available functional information.

To better understand how normalization of the CPTAC dataset using ProtMapper might add biological insight we chose a site found in the CPTAC dataset, IRS2-S560, that is found in PhosphoSitePlus but has no functional annotations in any database. All three text mining systems found multiple publications in which IRS2 is phosphorylated at residue “S556” by the kinase PLK1 (Chen et al., 2015; Mao et al., 2016, 2018). For example, Chen et al. described the potential regulatory significance of this phosphorylation event in human cell lines as follows: *“…Plk1-dependent phosphorylation of IRS2-S556 inhibits mitotic exit, partially through reduced AKT activity.”* Although the human IRS2 protein reference sequence does not have a serine at S556, ProtMapper correctly identifies this as a site on *murine* IRS2 that corresponds to site S560 on human IRS2 (both human IRS2 S560 and murine IRS2 S556 are known to be phosphorylated and are in the same site group in

PhosphoSitePlus). Although Chen et al. used the human HEK239T and HeLa cell lines for functional studies, their materials and methods section reveals that they raised an antibody against a recombinant fusion between the murine IRS2 protein and GST, then used the antibody to assay human cell extracts (Chen et al., 2015) — potentially explaining the use of a mouse site position for a study in human cell lines. Subsequent articles citing Chen et al. also incorrectly referenced the non-human site position S556 for human studies, for example, in the context of human pancreatic cancer cell lines (Mao et al., 2016, 2018). This example illustrates how a subtle inconsistency in a paper using both murine and human proteins can generate an invalid residue assignment that is then propagated to subsequent publications. The error has potential biological implications — both IRS2 (Tan et al., 2010, p. 5) and PLK1 (R. Zhang et al., 2015) have been reported as relevant to ovarian cancer treatment or prognosis, and IRS2 carries signals from insulin and insulin-like growth factor to the PI3K/AKT signaling pathway, which also plays a role in ovarian cancer (Ediriweera et al., 2019). This example highlights how text mining tools help link structured information about phosphorylation events to specific results in the primary literature — making it possible to evaluate the provenance, context-specificity and significance of PTM annotations.

## Discussion

This paper describes a method, implemented as Python software, for increasing the breadth and depth of functional information about human PTMs, with a focus on phosphorylation. We find that literature and phosphoproteomic datasets contain many references to non-canonical sites of modification, thereby impeding access to available functional information on these sites. Normalizing positions of PTMs to canonical reference sequences greatly facilitates assembly of site information from multiple databases and from text mining tools. We show that ProtMapper, coupled with the INDRA knowledge assembly system, can be used to create a corpus of functional phosphosite information that is 2.6-fold larger than the current standard (PhosphoSitePlus). Use of this corpus nearly doubles the fraction of sites with known regulators in datasets generated by CPTAC. To our knowledge the corpus we have assembled represents the most comprehensive source of literature-derived information about phosphosite regulation currently available. While the incorporation of text-mined regulatory information into the corpus introduces the possibility of technical errors, the use of INDRA’s belief scores allows the impact of these errors to be controlled in downstream applications.

Our analysis of phosphosite information extracted from databases and mined from the literature reveals that inconsistencies in site numbering are common in both sources: database curators and machine reading systems are misled by inconsistent references to amino acid sites in the literature. These errors appear to originate from historical bias in early functional studies involving a protein isoform that is no longer considered canonical, or to non-human species (particularly mouse or rat) if experimental materials from those species are used in the study (as illustrated above in the example of raising an antibody against mouse IRS2-S556 to study human IRS2-S560). Moreover, antibody manufacturers often list non-reference site positions on their product datasheets and these too are propagated in the literature. Resolving these inconsistencies is challenging and time-consuming for human curators. As a result, some databases, NCI-PID for example, contain a high proportion of invalid site annotations. By automating the process of site normalization ProtMapper has the potential to streamline the maintenance of phosphoproteomic and pathway databases (e.g., SIGNOR and Reactome) and thereby improve their scope and accuracy.

A substantial proportion (33-39% depending on the reader) of phosphosite annotations extracted by machine readers are technically correct with respect to reading accuracy but invalid with respect to a reference sequence for the protein described in the text (**Table 1**). Without ProtMapper, the information on these sites would simply be discarded from further analysis, but with ProtMapper we found that roughly a third of these sites could be “rescued” by mapping them to canonical positions. In our manual evaluation of 100 text mined sites, we found that ProtMapper made these mappings with 95% precision, a figure that can be increased still further by disabling the off-by-one rule used to correct for cleavage of the initiator methionine. The high precision and recall metrics show that ProtMapper reliably identifies valid information from text mining that would otherwise be indistinguishable from reading errors, while introducing very few false positives. Automated site normalization therefore plays an essential role in reliably assembling text mined information about PTMs.

We found that text mining systems freely available in the public domain can curate a substantial amount of information on phosphosite regulation that is currently “lost” to systematic analysis. This includes not only information similar to what is generally found in curated databases (direct interactions between individual kinases and substrates) but also regulation specified at the level of kinase families and indirect regulation by ligands, receptors, and upstream kinases. These types of functional annotations can be particularly valuable for phosphoproteomic data analysis because they allow higher-level patterns of regulation (e.g., by kinase families or signaling pathways) to be identified by straightforward enrichment analysis. Co-assembling text mining results with curated databases therefore extends both the size and diversity of available corpora on phosphosite regulation.

One limitation of ProtMapper’s current approach is that it uses human proteins as the point of normalization and is therefore not immediately useful for phosphoproteomic studies of distantly related species (e.g., *Drosophila* or yeast). However, ProtMapper is in principle species-agnostic and can therefore be extended to additional organisms in the future as the underlying experimental and literature-based information becomes available. ProtMapper can also be extended to other types of PTMs such as ubiquitination, methylation, and acetylation using a conceptually similar approach. A second limitation of ProtMapper is that its approach to mapping relies on prioritized set of predefined matching rules (**Figure 2C**) rather than a probabilistic approach that might, in principle, resolve conflicts between multiple possible mappings (e.g., an invalid site caused by an erroneous use of a mouse residue number or, alternatively, initiator methionine cleavage, which would result in different mappings). Despite this, our evaluations show that the current implementation of ProtMapper results in high precision mappings (**Table 3**).

In conclusion, this paper illustrates the need for new tools to effectively aggregate data on PTMs and their regulators at proteome scale. Fortunately, relatively simple tools such as ProtMapper can have a substantial positive impact. However, fully resolving inconsistencies and ambiguities in functional descriptions of proteins will require the involvement not only of database curators but also authors, editors and antibody manufacturers. Adoption of standard names for genes and canonical residue numbers will improve reproducibility, reusability and machine readability. ProtMapper could be used, for example, to check all phosphoprotein sites in submitted papers prior to their publication and to normalize all of the non-canonical annotations in vendor catalogs. ProtMapper is aimed at addressing only a handful of the problems associated with curating and integrating proteomics data and the field will benefit as the ecosystem of similar open-source software expands.

## Materials and Methods

### Information Sources

- PhosphoSitePlus: Downloaded from the PhosphoSitePlus website (https://www.phosphosite.org) on June 3rd 2022. Kinase-substrate annotations were obtained by processing the Kinase_substrate.owl BioPax file with the INDRA BioPAX processor into INDRA Statements. The file Phosphorylation_site_dataset.tsv was used by the ProtMapper for site mappings. License: CC BY-NC-SA 3.0. See also license information at https://www.phosphosite.org/staticDownloads.
- HPRD: Obtained from http://www.hprd.org/RELEASE9/HPRD_FLAT_FILES_041310.tar.gz and processed to INDRA Statements by the INDRA HPRD Processor.
- SIGNOR: Interactions obtained from https://signor.uniroma2.it/ on June 3rd 2022 and processed to INDRA Statements by the INDRA SIGNOR Processor. License: CC BY-SA 4.0.
- BEL Large Corpus: Downloaded from https://arty.scai.fraunhofer.de/artifactory/bel/knowledge/large_corpus/large_corpus-20170611.bel.
- Reactome: Obtained from Pathway Commons v12 and processed with the INDRA BioPAX processor. License: CC0.
- NCI-PID: Obtained from Pathway Commons v12 and processed with the INDRA BioPAX
processor.
- Reach: Software obtained from https://github.com/clulab/reach and used to process a text corpus including MEDLINE abstracts and full-text articles from the PubMed Central Open Access Subset, the PubMed Central Author’s Manuscript Collection, and others obtained via the Elsevier text and data mining API (https://dev.elsevier.com/).
- Sparser: Executable software image obtained from the Sparser developers and used to process the same text corpus as Reach.
- RLIMS-P: Text mining results for PubMed Central full-text articles and MEDLINE abstracts obtained via download from the iTextMine service (Ren et al., 2018) at https://hershey.dbi.udel.edu/textmining/export/, and processed to INDRA Statements using the INDRA RLIMS-P Processor. License: CC BY-NC-SA 4.0.
- UniProt: Protein identifiers, annotations, sequences and mappings to RefSeq identifiers were obtained from the UniProt website, https://www.uniprot.org. Specific download procedures are implemented in protmapper.resources.
- RefSeq: Protein sequences obtained from ftp://ftp.ncbi.nlm.nih.gov/genomes/Homo_sapiens/protein/protein.fa.gz.

### Mass Spectrometry Data Sources

CPTAC phosphoproteomic data were downloaded from the NCI Proteomic Data Commons (https://pdc.cancer.gov/pdc/). Breast cancer data was extracted from the CPTAC Prospective Breast BI Phosphoproteome study (https://pdc.cancer.gov/pdc/study/PDC000121) using the CPTAC2_Breast_Prospective_Collection_BI_Phosphoproteome.phosphosite.tmt10.tsv data file.

Ovarian cancer data was extracted from the TCGA Ovarian PNNL Phosphoproteome Velos Qexactive study (https://pdc.cancer.gov/pdc/study/PDC000115) using the

TCGA_Ovarian_PNNL_Phosphoproteome.phosphosite.itraq.tsv data file.

### Manual site curation

Inconsistent site positions in the NCI-PID database were manually examined and matched to site positions in the protein reference sequence. Internet searches of genes and their inconsistent site positions frequently identified both the source of the error (e.g., incorrect site position listed by antibody vendor) and the corresponding sites in the reference sequence. Incorrect sites were prioritized for curation by their frequency of appearance in BioPAX reactions. The resulting table listing incorrect sites, their reference positions, and a brief description of the source of the inconsistency is contained in the GitHub repository for the ProtMapper at: https://github.com/indralab/protmapper/blob/master/protmapper/curated_site_map.csv.

### Curating the accuracy of site normalization

Manual curation of site mappings for machine reading-derived sites were based on two criteria: 1) whether the site extracted by the reader was supported by the corresponding sentence in the source publication, and 2) whether the reference site returned by the ProtMapper was the correct one based on the context of the original sentence and publication. The criteria for scoring each site are summarized in Table 6.

**Table 6.**
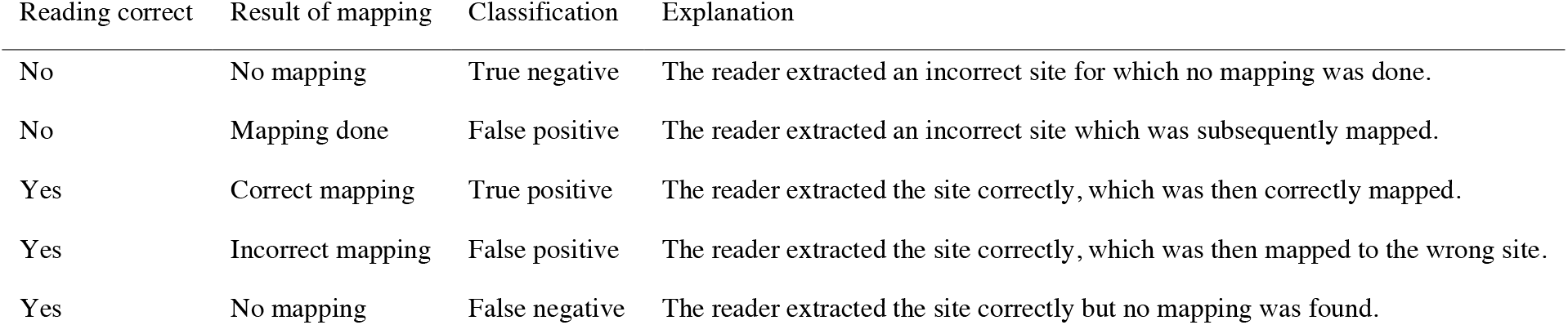
Curation categories for invalid sites extracted from the literature by machine reading and then mapped to human reference.

## Acknowledgements

Funding for this work was provided by the Defense Advanced Research Projects Agency under awards W911NF-14-1-0397, W911NF018-1-0124, and W911NF-20-1-0255, and by NIH grants U24-DK116204 and U54-CA225088. Data used in this publication were generated by the Clinical Proteomic Tumor Analysis Consortium (NCI/NIH). We thank Juliann Tefft for helpful comments on the manuscript.

## Availability

ProtMapper is implemented in Python and is available under a BSD 2-clause open-source license from GitHub at https://github.com/indralab/protmapper. Documentation is hosted at

https://protmapper.readthedocs.io. Code used to assemble the annotation corpus and generate the results in this paper is on GitHub at https://github.com/indralab/protmapper_paper. The assembled corpus of annotations is available under a CC-BY-NC-SA license (per the Share-alike requirements of constituent sources PhosphoSitePlus, RLIMS-P and SIGNOR) and is available as Supplementary Data with this paper.

## Author contributions

JAB and BMG conceived and implemented the software, and performed the analysis. JAB, BMG and PKS wrote the paper.

