## Supplementary material for "Assembling a corpus of phosphoproteomic annotations using ProtMapper to normalize site information from databases and text mining": LICENSE.txt

The following licenses apply to knowledge from sources listed in exports.csv and evidence.csv.

PhosphoSitePlus (psp): PSP is licensed by Cell Signaling Technology (CST) for
non-commercial use under a Creative Commons
Attribution-NonCommercial-ShareAlike 3.0 Unported License and the further terms
listed below. See https://www.phosphosite.org/staticDownloads for more details.

RLIMS-P / iPTMnet (rlimsp): The iPTMnet data are freely searchable and
browsable from the website. However, the iPTMnet database is protected under
the CC BY-NC-SA 4.0 (Attribution-NonCommercial-ShareAlike 4.0 International)
license. See https://research.bioinformatics.udel.edu/iptmnet/license for more
details
