## Supplementary material for "Assembling a corpus of phosphoproteomic annotations using ProtMapper to normalize site information from databases and text mining": README.txt

The export consists of two related tables: export.csv and evidence.csv that
are linked via the ID column in each table. Note that the IDs in export.csv
can map to several rows in evidence.csv since one annotated site
can have evidence from multiple sources.

Columns in export.csv:
- ID: the identifier of the site annotation
- CTRL_NS: the namespace of the controller with the following values:
(HGNC: Human Gene Nomenclarure Consortium, UP: UniProt, FPLX: FamPlex)
- CTRL_ID: the identifier of the controller within the given name space
- CTRL_GENE_NAME: the official gene symbol corresponding to the controller,
when available
- CTRL_IS_KINASE: True of False, depending on whether the controller is known
to be a human kinase or not
- TARGET_UP_ID: the UniProt identifier of the target protein
- TARGET_GENE_NAME: the official gene symbol corresponding to the target,
when available
- TARGET_RES: the residue of the final (valid or mapped) site on the target
- TARGET_POS: the position of the final (valid or mapped) site on the target
- SOURCES: a comma-separated, sorted list of sources from which there is evidence
for the given annotation
- BELIEF: a score between 0 and 1 representing confidence in the correctness
of the annotation, as calculated by INDRA. All annotations directly or
indirectly supported by curated databases have beliefs set to 1.

Columns in evidences.csv:
- ID: the identifier of the site annotation to which the evidence corresponds
(multiple rows can have the same ID if there are multiple evidences for the
same site annotation)
- SOURCE: the source of the evidence (either a database or a machine reading
system)
- PMID: the PubMed identifier of the paper from which the site was extracted,
when available. Note that for technical reasons, the current export is missing
PMIDs for a subset of evidences from the REACH and Sparser systems. In many
cases, the original article can be found by searching for the TEXT entry on the
web.
- DBID: the identifier of the database entry from which the site was extracted,
when available
- TEXT: the evidence sentence corresponding to the site
- DESCRIPTION: this field is either "VALID" meaning the site was valid to
being with, or another code representing the mapping operation that was done
to obtain the site
- ORIG_UP_ID: the original UniProt ID of the protein corresponding to the site
- ORIG_RES: the original residue of the site
- ORIG_POS: the original position of the site
- MAPPED_UP_ID: the mapped UniProt ID of the protein corresponding to the site
- MAPPED_RES: the mapped residue of the site
- MAPPED_POS: the mapped position of the site
